# DNA metabarcoding and environmental DNA across Brazilian biomes: trends, challenges, and conservation applications

**DOI:** 10.64898/2025.12.11.693676

**Authors:** Julia Terra Souza Torres, Ingrid Bunholi, Naiara Guimarães Sales, Rodrigo R. Domingues

## Abstract

Brazil harbors the greatest biodiversity on the planet; however, the application of emerging technologies to enhance biodiversity assessment and monitoring remains limited. In recent years, DNA metabarcoding and environmental DNA (eDNA) surveys have revolutionized the assessment of biodiversity, offering new tools to evaluate ecological and conservation trends and patterns worldwide. Yet, their application in Brazil is still in its early stages, underscoring the paradox of a megadiverse country with limited adoption of cutting-edge tools for monitoring its natural wealth. This review provides a comprehensive synthesis of the application of DNA metabarcoding and eDNA analyses across different ecosystems in Brazil, integrating findings from 47 studies published between 2015 and 2025. Trends in ecosystem types, taxonomic groups, molecular markers, geographic distribution, and patterns of authorship and funding were analyzed. Most studies focused on terrestrial (42.5%) and freshwater (34%) environments, followed by marine (19.1%) and estuarine (4.2%) systems. DNA metabarcoding predominated in terrestrial studies (52.1%), while eDNA was more frequent in freshwater, marine, and estuarine environments. The mitochondrial 12S marker was the most widely used, primarily for detecting fish and amphibians. We observed a predominance of Brazilian first and senior authors, with weak signs of parachute science. However, challenges persist, including limited access to funding and underrepresentation of some biomes and taxonomic groups. Our findings highlight a growing national capacity in the use of DNA metabarcoding and eDNA research while underscoring the need for strategic investments in infrastructure, database development, and equitable scientific collaboration.

## 1. Introduction

Biodiversity plays a fundamental role in the balance and functionality of ecosystems, holding ecological, economic, and social importance (Mace et al., 2018; Hong et al., 2022). In the past few decades, the mounting biodiversity loss has become one of the most critical environmental problems, with species and local extinction records at rates much faster than we can afford to study and monitor (Ceballos et al., 2015). During the 20th century, biodiversity declined by an estimated 2 to 11%, based on various indicators worldwide (Pereira et al., 2024). Despite decades of accumulated evidence, the trajectory of biodiversity in the Anthropocene remains uncertain, with attempts at synthesis yielding mixed and debated results (Keck et al., 2025), which underscore the need for innovative approaches to assess biological diversity.

Monitoring species provides essential insights into their presence, distribution, and ecological and conservation roles. It also enables the detection of endangered or invasive species and enhances understanding of ecosystem interactions (Lindenmayer et al., 2014; Sparrow et al., 2020). These insights are crucial for both biodiversity conservation and a deeper understanding of ecosystem functioning (Boulinier et al., 1998; Lepetz et al., 2009; Yates et al., 2023). Traditional sampling, whether invasive or non-invasive, involves direct collection or observation of organisms using techniques such as netting, trapping, and visual surveys. Terrestrial methods include line-transects, pitfall and camera traps, and acoustic monitoring (Braga-Pereira et al., 2021), while aquatic systems use fisheries-dependent data, underwater visual censuses, baited remote video systems, and remotely operated vehicles (Whitmarsh et al., 2017; Oka et al., 2021).

The effectiveness of sampling techniques varies by taxonomic group, from elusive mammals that are hard to detect (Burns et al., 2017) to species-rich groups like fish and invertebrates with challenges in monitoring exacerbated by cryptic taxa (Hending et al., 2025). Most long-standing approaches rely on morphological traits, which may be influenced by phenotypic plasticity, life-history variation, and observer expertise, potentially causing misidentifications, especially for cryptic or degraded specimens (Goldberg et al., 2011; Dejean et al., 2012; Jörger & Schrödl, 2013). To complement morphological species identification, DNA barcoding was developed as a molecular tool that uses a standardized DNA region for species identification (Hebert et al., 2003). Although initially limited to identifying one individual at a time (Hebert & Gregory, 2005), the relatively new expanding (ca. 15 years) integration of barcoding with high-throughput sequencing (HTS) technologies enabled the emergence of more comprehensive approaches such as DNA metabarcoding, which allows for the simultaneous detection of multiple taxa in complex biological mixtures (Hering et al., 2018).

DNA metabarcoding has largely contributed to the understanding of community composition (e.g., analyses of bulked samples; Lynggaard et al., 2020; Martins et al., 2025) and species interactions, enabling simultaneous detection of multiple taxa from complex mixtures. Applied to feces and gut contents, it delivers precise diet characterization, including cryptic, soft-bodied, and highly digested prey, revealing broader prey spectra and plant use than observational or morphological methods (de Queiroz et al., 2024; Rosa et al., 2024). DNA metabarcoding has also improved our understanding of fish habitat use and recruitment dynamics, pinpointing spawning and nursery areas and clarifying species-specific settlement patterns (Nobile et al., 2019; Silva et al., 2023). These data resolve trophic niches and seasonal resource partitioning among sympatric species and, when aggregated, allow reconstruction of multi-trophic food webs (e.g., predator–prey), strengthening network metrics and informing conservation of keystone species, interaction diversity, and ecosystem functioning.

The use of DNA metabarcoding coupled with environmental DNA (i.e., genetic material obtained directly from environmental samples, such as soil, water, or air, without the need to capture or visually identify the organisms, Taberlet et al., 2012) represented a major advancement in biodiversity monitoring. The eDNA metabarcoding merged as a powerful and non-invasive approach not only for biodiversity monitoring but also for exploring ecological interactions, community structure, functional diversity, and biogeographic patterns (Creer et al., 2016; Hering et al., 2018; Yates et al., 2023). For example, the ’Fun-eDNA’ approach introduced by Cantera et al. (2025) enables the characterization of functional diversity by assigning ecological traits to taxa identified through eDNA metabarcoding. This method provides a comprehensive perspective on ecosystem functioning and allows researchers to infer the functional roles of species within communities, such as their positions in food webs or their contributions to nutrient cycling, thereby enhancing our understanding of biodiversity patterns and ecosystem processes (Cantera et al., 2025).

Although the application of eDNA has increased exponentially over time, this growth is not geographically uniform, with a marked prevalence of studies in the Northern Hemisphere (Bunholi et al., 2023; Takahashi et al., 2023; Broadhurst et al., 2025). Such disparity is particularly evident in low and middle-income countries, where limited research funding and restricted access to molecular infrastructure hinder broader adoption (Polejack & Coelho, 2021). As highlighted by Bunholi et al. (2023), South America and Africa together represent less than 10% of global eDNA-based biodiversity assessments, revealing a persistent geographic bias that may obscure biodiversity hotspots and critical ecosystem services in these underrepresented regions (Heyden, 2022).

Brazil is globally recognized as the most biodiverse country, classified as a megadiverse nation due to its extensive terrestrial and aquatic ecosystems (Forzza et al., 2012; ICMBio, 2018; Brasil, 2024). It harbors six terrestrial biomes and the highest known numbers of plant (Forzza et al., 2012), amphibian (Vié et al., 2009), primate (Brito et al., 2009; Sampaio et al., 2017), and freshwater fish species (Albert & Reis, 2011; Dagosta & de Pinna, 2019), along with high endemism rates, especially among reptiles and amphibians (Mittermeier & Goettsch Mittermeier, 1997; Costa & Bérnils, 2014). Brazil also supports significant marine biodiversity and priority conservation areas along its 7,491 km coastline, and is recognized as an important area of marine endemism (Miloslavich et al. 2011). In this context, DNA-based tools like eDNA and DNA metabarcoding have emerged as powerful and cost-effective alternatives to species identification, enabling biodiversity monitoring surveys, the exploration of complex ecological processes, and improving our capacity to inform management and conservation strategies across diverse environments (Thomsen and Willerslev, 2015; Dugal et al., 2022; Diniz-Filho et al., 2024).

Given Brazil’s exceptional biodiversity, our ultimate goal is to demonstrate how eDNA and DNA metabarcoding can be applied to community monitoring, and ecosystems and species conservation in a megadiverse country. We synthesize studies that deploy these tools to address ecological questions, from biodiversity assessment to trophic dynamics, across Brazilian ecosystems, and we summarize (i) the geographic and (ii) taxonomic coverage, (iii) the principal molecular markers used, (iv) the taxonomic units analyzed, and (v) the breadth of ecological applications. We also examine evidence of parachute science and outline practical, conservation-focused directions for future eDNA/metabarcoding surveys in Brazil.

## 2. Methods

A systematic literature survey was conducted in December 2024 and additionally in May 2025 on Web of Science, Scopus, and Google Scholar databases, using Boolean operators. The following query was used: “(ALL= Brazil AND Metabarcoding OR Brazil AND Metabarcode) AND (ALL= Brazil AND eDNA OR Brazil AND environmental DNA)”. In this study, the term DNA metabarcoding means an approach for the detection of multiple taxa from complex mixtures (e.g., tissue bulk, feces, gut content), whereas eDNA metabarcoding (hereafter eDNA) means the detection of multiple taxa from environmental samples (e.g., water, soil, sediment). No studies using eDNA in combination with other molecular methods (e.g., qPCR or ddPCR) were identified. Hereafter, all studies were analyzed together and treated as DNA metabarcoding and eDNA studies, divided into different categories.

A first filter was applied to exclude reviews and studies targeting viruses, bacteria, and fungi. A second filter applied retained only studies conducted in Brazil. After applying these filters, 208 papers were retained. A manual inspection was carried out to remove preprints, reviews, papers using other molecular techniques (e. g., DNA barcoding) or not focused on Brazilian ecosystems. Finally, 47 papers were selected, scanned, and divided into four categories according to environment: marine, terrestrial, freshwater, and estuarine. Among the 47 selected publications, 23 studies applied DNA metabarcoding to non-eDNA samples such as bulk organismal samples, feces, stomach content, or tissue, whereas 24 used eDNA samples. A metadata file containing all metrics collected from all papers was created to generate the figures and statistical metrics (Supplementary Information S1). These metrics included: authors, title, year of publication, study applicability (e.g., conservation, trophic interactions, monitoring, etc), sample (water, sediment, tissue, etc), environment (terrestrial, marine, freshwater or estuarine), region (state and biome), target organism (e.g., fish, invertebrate, amphibian, arthropod non-target, etc), extraction method, targeted genetic marker, and reported taxonomic unit (ASV and/or OTU). For studies with multiple target genetic markers, ecosystems, or target organisms, we treated each one independently. To classify the studies across ecosystems, we considered the target organisms as the primary criterion rather than the sample type. In most cases, the sample origin and focal organisms matched (e.g., water samples for aquatic organisms, soil samples for terrestrial taxa). Five exceptions were observed: one study used water samples to detect both aquatic and terrestrial mammals, which we classified as aquatic (Sales et al., 2019a), another used bromeliad water to assess terrestrial amphibians (Lopes et al., 2021), and three studies collected stream water, also to investigate terrestrial amphibians. These latter four studies were classified as terrestrial (Lopes et al., 2017; Lopes et al., 2020b; Ernetti et al., 2024).

To assess the prevalence of parachute science among the selected studies, we examined both authorship composition and sources of funding. Parachute science refers to research primarily led by scientists from the Global North, conducted in countries of the Global South without the involvement or meaningful collaboration of local researchers or stakeholders (Asase et al., 2021; Stefanoudis et al., 2021; de Vos & Schwartz, 2022; Heyden, 2022). This analysis aimed to identify cases of absent or minimal local participation, thereby highlighting imbalances in scientific collaboration and authorship.

The studies were categorized according to: (i) whether first and/or senior authors were affiliated with foreign institutions; (ii) the presence of foreign-affiliated authors elsewhere in the author list; (iii) the absence of any foreign representation; (iv) exclusive foreign funding; and (v) combined Brazilian and foreign financial support. In cases where Brazilian researchers were affiliated with institutions abroad, the nationality of the authors was taken into account to more accurately reflect local participation.

All analyses were performed on the statistical software R version 4.4.3 (R Core Team, 2025).

## 3. Results and Discussion

The 47 papers retained after the filtering process were published across 33 scientific journals. The most frequently represented journals were Environmental DNA (n = 4; 8.7%), Scientific Reports (n = 4; 8.5%), PeerJ (n = 4; 8.5%), and Ecology and Evolution (n = 3; 6.4%). All selected studies were published between 2015 and 2025, showing interannual variation and an average rate of 4.6 studies per year, with a peak in 2023 when 11 studies were published (Figure 1). This increase in publications over the past decade aligns with the growing adoption of DNA metabarcoding and/or eDNA-based tools in biodiversity research (e.g., Minamoto, 2022; Nordstrom et al., 2022; Schenekar, 2023).

**Figure 1.**
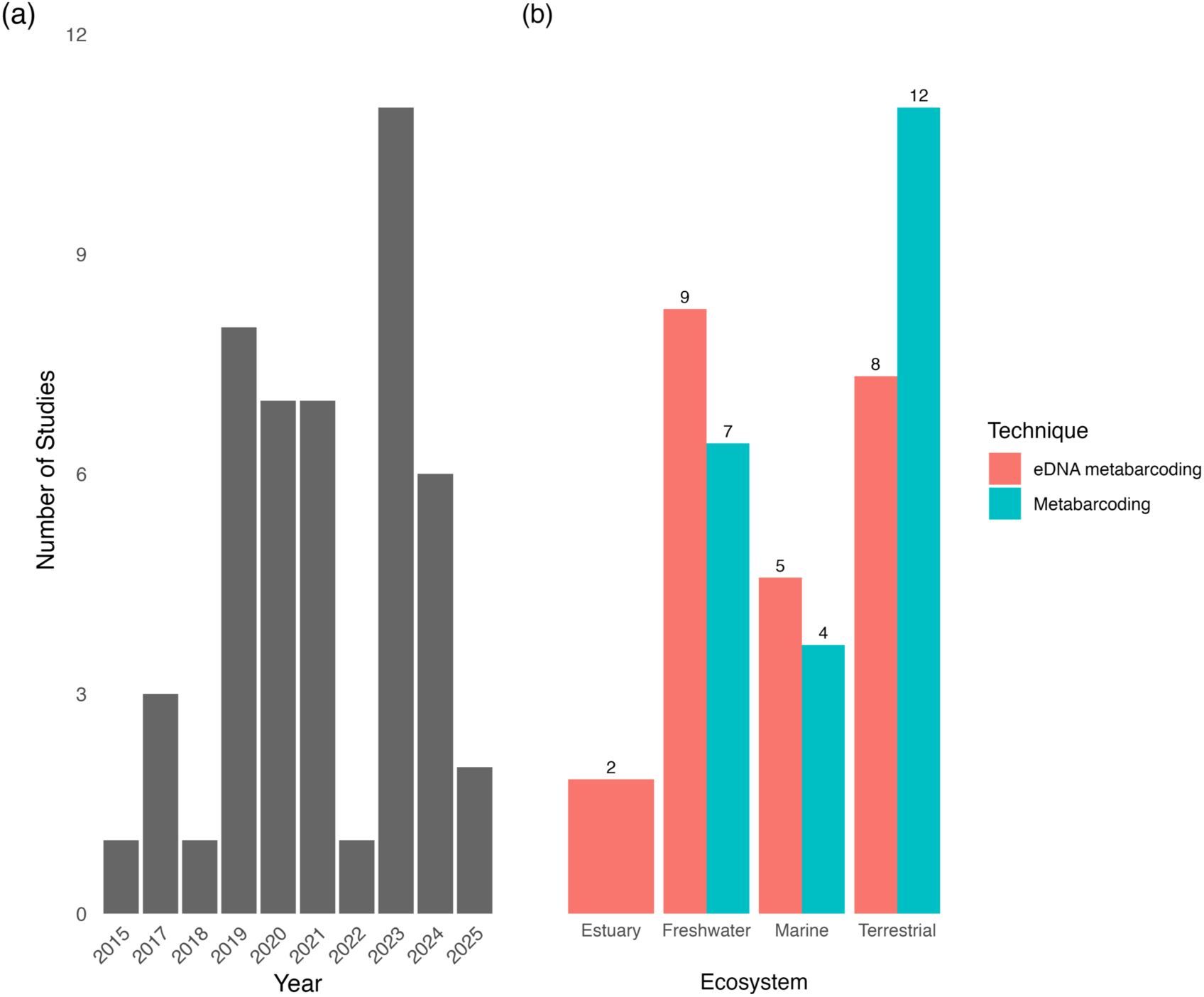
Number of published studies using metabarcoding and eDNA metabarcoding in Brazil between 2015 and 2025. (a) Annual number of studies using either technique. (b) Total number of studies using each technique per ecosystem (Estuarine, Freshwater, Marine, Terrestrial). Values above bars represent the exact number of studies.

Regarding applicability, the 47 selected studies were assigned to different categories, reflecting the broad scope of research. The five most frequent applicability categories were biodiversity monitoring (n = 22; 46.8%), conservation (n = 11; 23.4%), trophic interactions (n = 8; 17%), species distribution (n = 4; 8.5%), and biological invasion (n = 2; 4.2%). This categorization highlights the main direction of DNA metabarcoding and/or eDNA research in Brazil.

### 3.1 Studied ecosystems

Among the analyzed papers, 20 were conducted in terrestrial environments (42.5%), of which 12 (60%) applied DNA metabarcoding and eight (40%) used eDNA. Freshwater environments accounted for 16 papers (34%), including seven (43.7%) based on DNA metabarcoding and nine (56.2%) on eDNA. In marine environments, nine papers (19.1%) were identified, four (44.4%) using DNA metabarcoding and five (55.6%) using eDNA, while the two papers in estuarine environments (4.2%) employed eDNA exclusively (Figure 1).

Regarding eDNA, Cowgill et al. (2025) demonstrated that terrestrial eDNA studies comprise less than 20% of global efforts in the field, with this proportion remaining relatively stable over the past decade, indicating a persistent underrepresentation of terrestrial systems. Similarly, in the Brazilian context, although nearly 50% of the reviewed studies were conducted in terrestrial environments, only eight of them employed eDNA. The remaining studies used DNA metabarcoding on non-environmental samples, highlighting that eDNA applications remain limited in terrestrial ecosystems. Furthermore, unlike the findings of Bunholi et al. (2023), whose global review of eDNA and eRNA studies in aquatic environments reported a predominance of marine research, our analysis indicates that freshwater systems have been more frequently investigated than marine ones in Brazil.

The first eDNA study for biodiversity assessment in Brazil was conducted by Lopes et al. (2015), who applied this approach to investigate the community composition of anuran species in the Atlantic Forest of São Paulo State. Most subsequent studies were conducted in the states of southeastern Brazil, including São Paulo (n = 11; 23.4%), Espírito Santo (n = 7; 14.9%), and Minas Gerais (n = 6; 12.7%). Studies from the northern Brazilian states such as Pará (n = 4; 8.7%), Amazonas (n = 5; 10.6%), and Rondônia (n = 3; 6.4%) collectively represent 25.5% of the total (Figure 2). This proportion is slightly higher than that of São Paulo alone, which stands out as Brazil’s most socioeconomically developed state. Despite the magnitude and ecological importance of the Amazon Rainforest, recent research has shown that northern Brazil receives the fewest scholarships and grants per square kilometer, and that the current federal budget is insufficient to cover large-scale research in the Amazon (Stegmann et al., 2024). This pattern reflects a broader global trend in which eDNA research is predominantly concentrated in economically developed countries and regions (Wang et al., 2021; Miya, 2022). Hirsch et al. (2024) further highlighted the concern that, if international standards are established predominantly by researchers from highly developed countries, they may become inaccessible or cost-prohibitive for those working in less-resourced (e.g., Global South) and remote regions.

**Figure 2.**
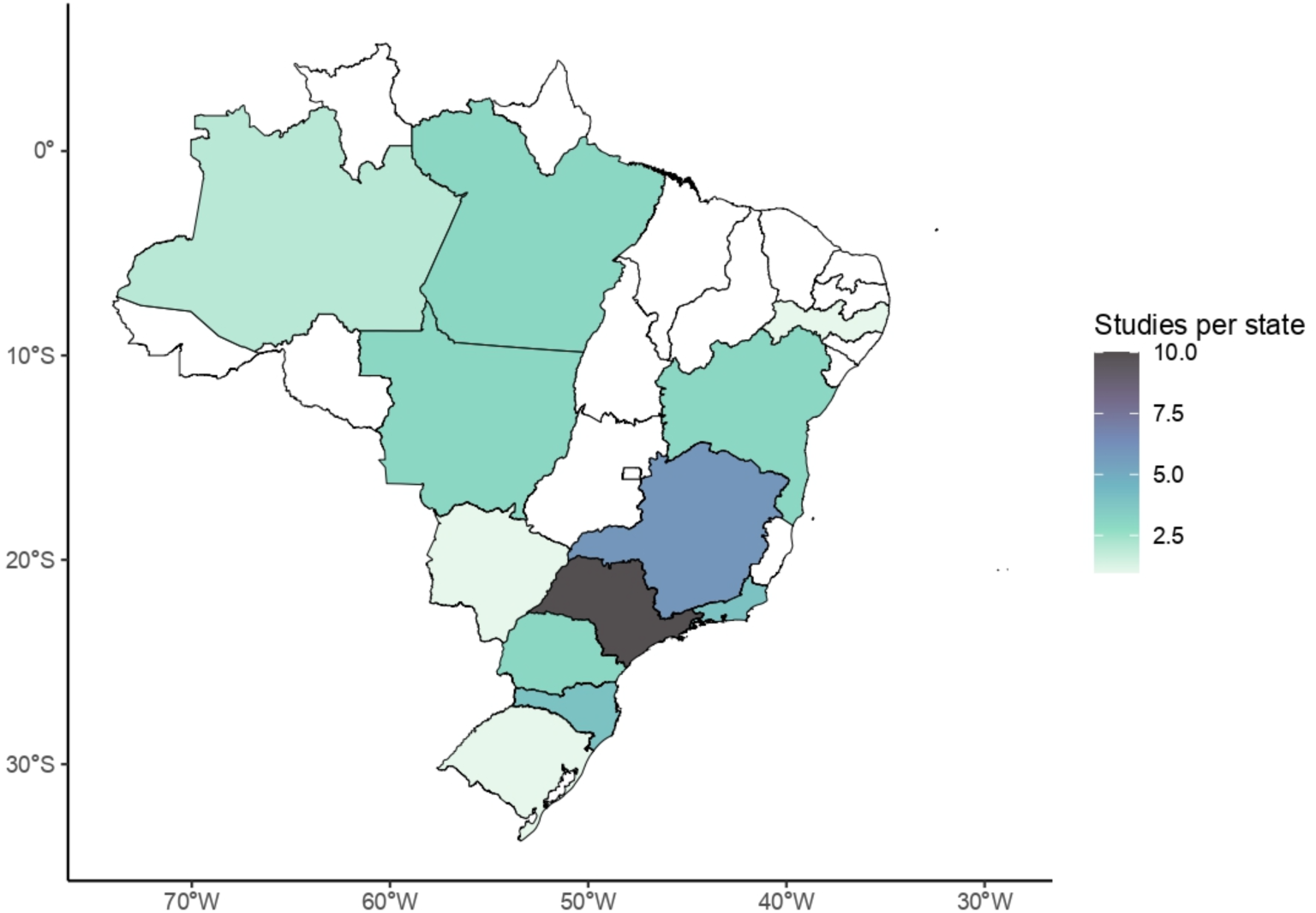
Number of studies that used environmental DNA (eDNA) and/or metabarcoding in each Brazilian state.

Among the twenty studies conducted in terrestrial environments, the main biomes investigated were the Atlantic Forest (n = 13; 65%) and the Amazon Rainforest (n = 8; 40%), followed by the Cerrado (n = 3; 15%) and Pantanal (n = 1; 5%) (Figure 3). Sixteen (n = 9 eDNA and n = 7 DNA metabarcoding) studies were conducted in freshwater ecosystems in Brazil, which were primarily focused on main rivers (n = 8; 53.3%) and river basins (n = 6; 40%), such as the São Francisco River Basin (n = 4; 26.7%), which spans multiple states across the country. Other important rivers investigated using DNA metabarcoding and/or eDNA techniques include the Jequitinhonha River (n = 3; 20%). Although other studies in the Amazon region have collected water from different rivers, only one specifically targeted the Amazon River itself, where water samples were collected to detect both aquatic and terrestrial mammals (Sales et al., 2019a). Reservoirs were another frequently studied environment in freshwater assessments, accounting for 33.3% (n = 6) of the studies.

**Figure 3.**
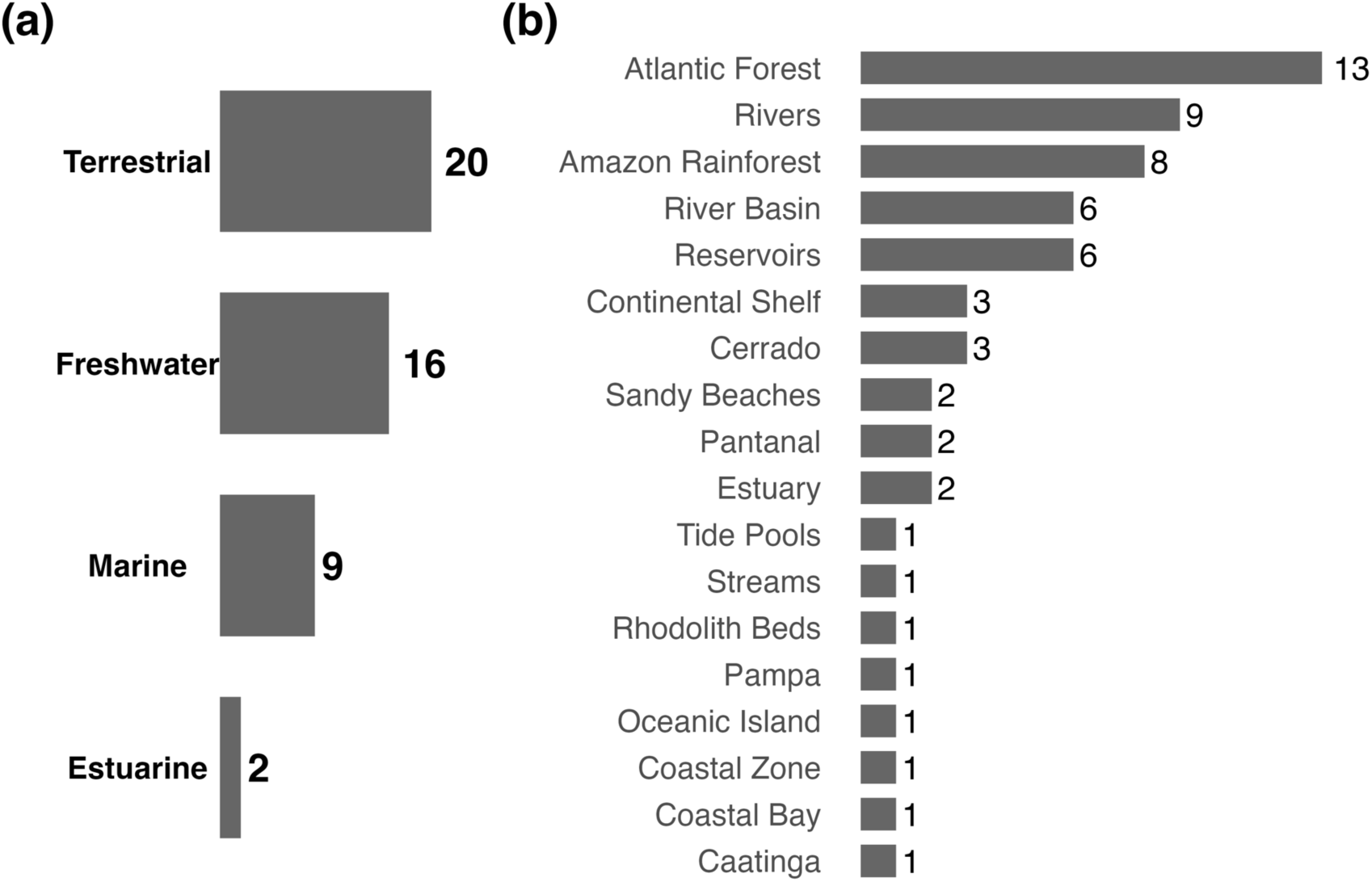
Distribution of sampling efforts across broad ecosystem types and environmental domains. (a) Number of studies conducted in each major ecosystem category: terrestrial, marine, freshwater, and estuarine. (b) Frequency of sampling across specific environmental domains.

In marine ecosystems, of the nine studies analyzed, most were conducted on the continental shelf (n = 4; 44.4%), primarily developing diet analysis through on-board monitoring of the sardine fishery along southeastern Brazil, spanning from Santa Catarina to Rio de Janeiro. Studies conducted in coastal environments (n = 3; 33.3%), focused on sandy beaches for benthic community surveys in Espírito Santo state and on coastal bays such as Araçá Bay, located on the northern coast of São Paulo State (Figure 3).

The only two studies conducted in estuarine ecosystems were carried out in the Espírito Santo state (Figure 3). Bernardino et al. (2019) assessed the ecological health through the analyses of benthic communities of the Rio Doce estuary 1.7 years (2017) after the initial impacts of the Samarco mining disaster. Subsequently, Coppo et al. (2023) investigated the same estuary to evaluate its condition both 1.7 years (2017) and 2.8 years (2018) after the release of mine tailings, aiming to understand the prolonged effects of sediment contamination on estuarine ecosystems.

### 3.2 Main molecular markers used

Across the 47 studies analyzed, nine distinct genetic markers were employed, with 12S (n = 19; 42.5%), COI (n = 14; 29.8%), 18S (n= 9; 17%), 16S (n= 7; 14.9%) and 28S (n= 3; 6.4%) representing the most frequently used (Figure 4). Eight studies (17%) applied a multimarker approach, enhancing taxonomic coverage and detection efficiency by combining markers with different specificity and resolution. For example, Ritter et al. (2019) employed a multimarker approach (COI, 16S, and 18S) to assess both prokaryotic and eukaryotic diversity and to evaluate the taxonomic coverage. Lines et al. (2023) used 16S to detect fish and marine mammals and COI for elasmobranchs. Three (60%) of the five most commonly used markers (12S, COI, 16S) are mitochondrial, which are typically preferred because of their high copy number per cell, maternal inheritance, lack of recombination, and greater persistence in degraded samples (Harrison et al., 2019). Nuclear markers (e.g., 18S, ITS2, 28S) were less used, as each cell typically contains only one nucleus with two copies per gene, compared to hundreds or thousands of mitochondrial DNA copies (Duarte, Simões, & Costa, 2023).

**Figure 4.**
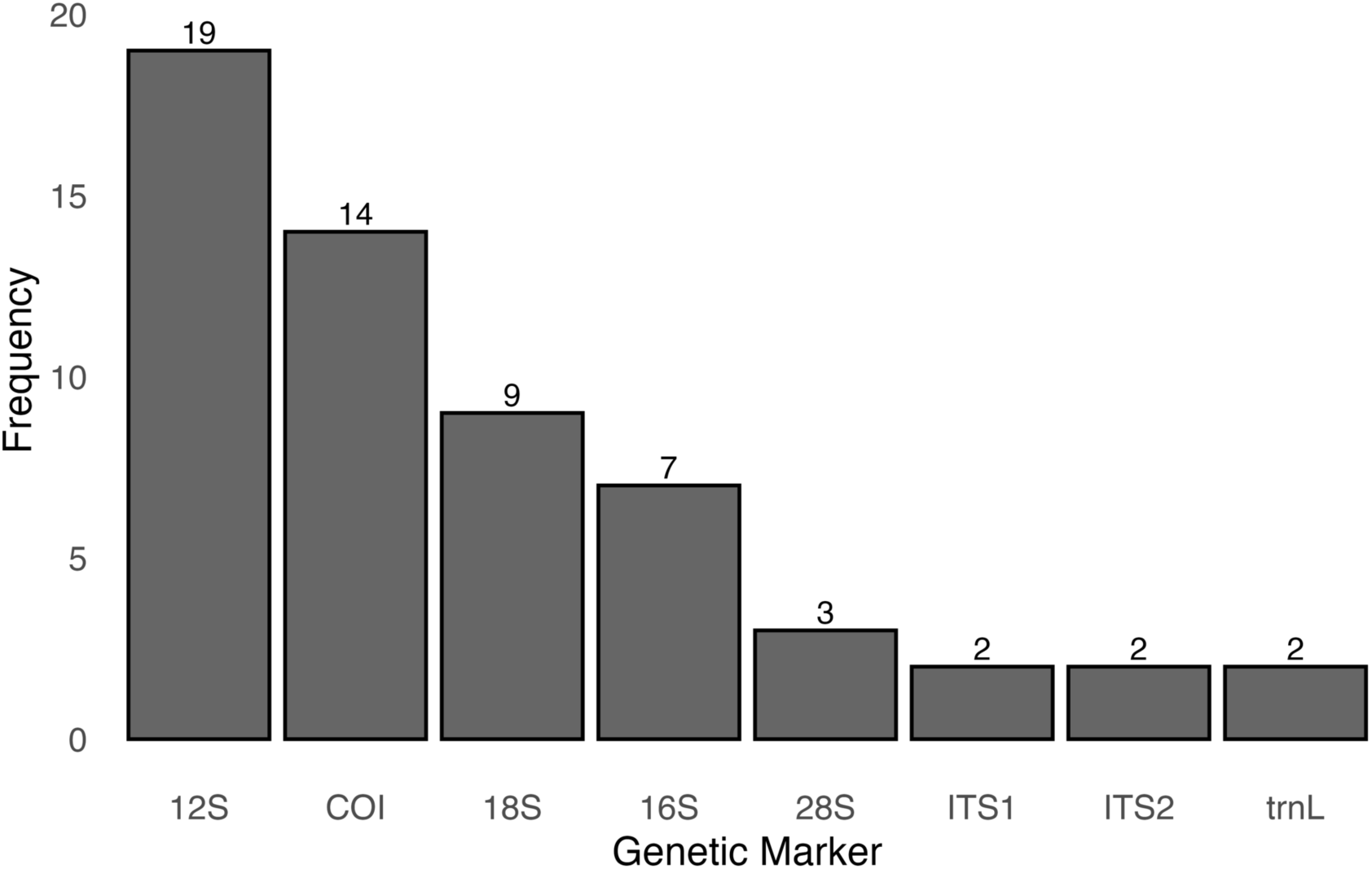
Frequency of molecular markers used in the eDNA and/or metabarcoding studies analyzed.

The 12S marker predominated in both terrestrial and freshwater environments (Figure 5a), mainly for detecting amphibians and fish (Figure 5b). In terrestrial systems, it was the exclusive choice for amphibians (Figure 5b), except in one study that combined 12S and 16S to detect mammals and herpetofauna (Lima et al., 2024). In freshwater and marine habitats, 12S was used mainly for fish (teleost and elasmobranchs) identification (Figure 5a, b). Only one study has applied the 12S marker for the identification of fish (elasmobranchs) in the marine environment (Cruz et al., 2023), and so far, there have been no applications for teleosts. This demonstrates a critical research gap, as expanding the use of molecular tools in marine conservation units could significantly enhance biodiversity monitoring and support more effective management strategies. The preference for 12S in vertebrate monitoring is due to its highly conserved regions of 20-30 base pairs flanked by hypervariable regions of ∼ 200 bp (Deagle et al. 2014, Valsecchi et al., 2020; Liu et al., 2023).

**Figure 5.**
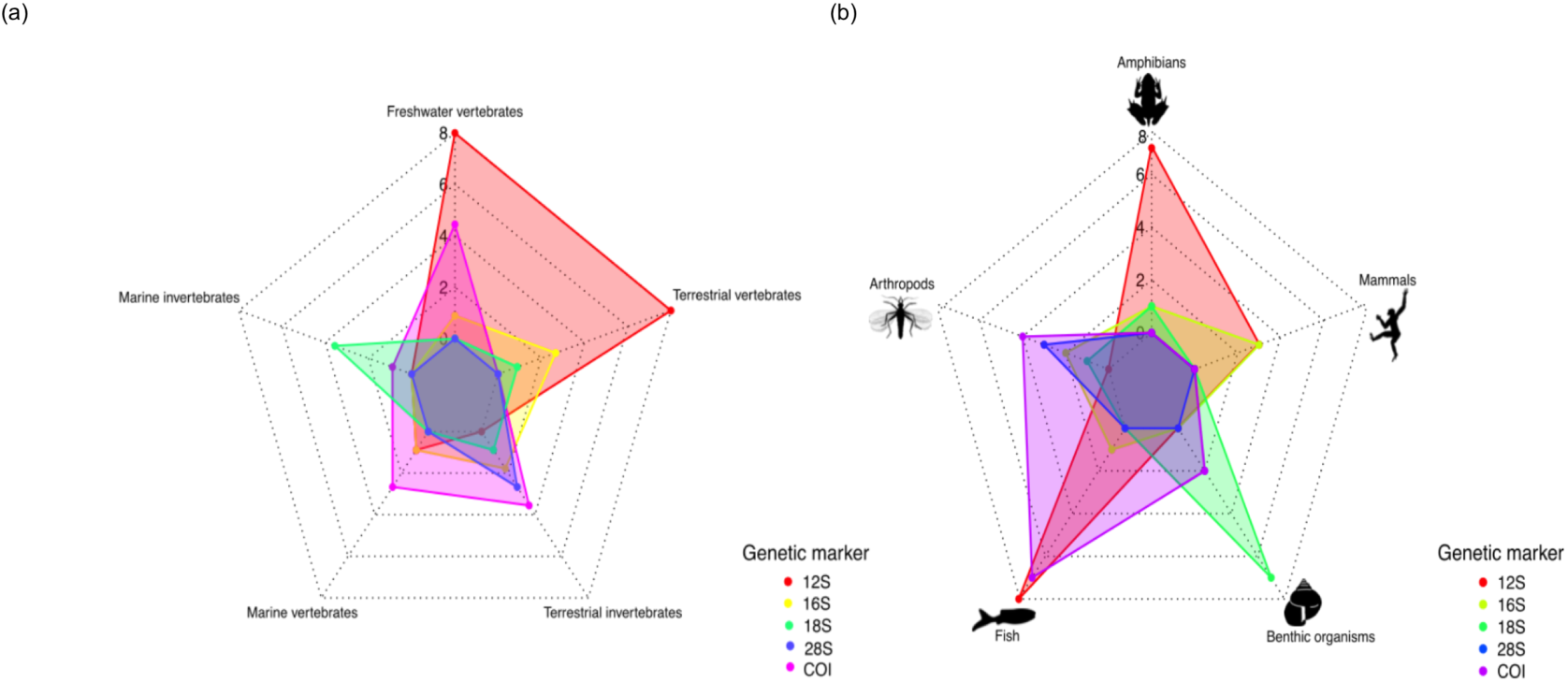
The five most used molecular markers across the five most targeted organismal groups studied by environmental DNA (eDNA) and metabarcoding in Brazil.

The COI gene was widely applied for arthropods in terrestrial environments and for fishes in marine and freshwater systems (Figure 5a, b). Its extensive use is supported by the availability of comprehensive reference databases, universality across metazoans, well-established protocols, and the high interspecific variability enables species-level resolution (Porter & Hajibabaei, 2018). However, primer degeneracy can reduce amplification efficiency for certain taxa, including some fish and mammals (Rennstam Rubbmark et al., 2018). For example, de Queiroz et al. (2024) found that most dietary items of a keystone sardine species were other fishes rather than crustaceans, contrary to expectations. This mismatch was likely due to the known difficulty of amplifying crustacean DNA with some COI primers (Zhan et al., 2014), underscoring the value of multimarker approaches to minimize false negatives and improve taxonomic recovery (Burian et al., 2023).

The 18S was used in eight studies (17%), and the sole molecular marker in estuarine research, both targeting benthic meiofaunal eukaryote communities (Bernardino et al., 2019; Coppo et al., 2023), and was also widely applied in marine environments for meiofaunal surveys (Figure 5a, b). Its broad taxonomic coverage and ability to detect small, morphologically cryptic organisms make it particularly suited for assessing benthic diversity (Ficetola et al., 2021; Liu et al., 2021). For example, de Faria et al. (2018) showed that 18S metabarcoding revealed greater meiofaunal diversity than morphological methods, while Coppo et al. (2024a, 2024b, 2025) documented high benthic diversity in rhodolith beds and intertidal zones along the coast of Espírito Santo, highlighting the role of habitat filtering and environmental drivers in shaping assemblages. In terrestrial systems, 18S appeared only in multimarker approaches (Lopes et al., 2020c; Ritter et al., 2019), often to complement more taxon-specific markers.

The 16S marker was applied mainly for detecting vertebrates in both terrestrial and aquatic environments, but appeared as the sole marker in only one case to detect arthropods (Figure 5a, b). Although 16S offers broad applicability and good resolution for certain taxa (Elbrecht et al., 2017; Sousa et al., 2019), Paula et al. (2022) reported that it failed to detect prey species in gut content analysis of arthropod epigeal, likely due to primer mismatches and preferential amplification of host DNA.

The 28S was reported in only three terrestrial studies (6.4%), all targeting mosquitoes using the D2 rDNA segment (Figure 5a, b). The D2 region was chosen for its higher taxonomic resolution and robust amplification from complex or degraded samples, overcoming common limitations of other markers, such as low taxonomic resolution, poor primer universality, and limited amplification success with degraded material, thereby offering a more practical and effective tool for ecological and vector monitoring (Beebe et al., 2018; Pedro et al., 2020).

Finally, the ITS2 marker was applied in only two terrestrial studies (4.3%) (Figure 4), targeting plants and pollen. Martins et al. (2023) used it to assess Neotropical bee foraging behavior, revealing a broader range of food sources than detected by conventional methods, while Vasconcelos et al. (2021) employed it to survey vascular plant diversity in Amazonian canga ecosystems. ITS2 is suitable for multispecies surveys due to its short amplicon size and ease of PCR standardization (Chen et al., 2010; Richardson et al., 2015; Gous et al., 2019). ITS1, also restricted to terrestrial studies, was used in combination with trnL and 16S to analyze subterranean rodent diets, enabling accurate identification of consumed plant species (Lopes et al., 2015, 2020a).

### 3.4 Main groups/ species studied

The majority of the studies (57,4%, n=27) targeted vertebrates, and 18 (38,3%) targeted invertebrates (Figure 6a). Fish were the most frequently studied group (n = 14; 29.8%), followed by arthropods (n = 9; 19.1%), amphibians (n = 7; 14.9%), benthic communities (n = 6; 19.1%), and mammals (n = 5; 10.6%) (Figure 6b). Only one study did not focus on a specific taxon (2.1%) (Ritter et al., 2019) (Figure 6b). All studies focused on animals, except one that investigated vascular plant biodiversity in the Amazon Rainforest (Figure 6). Vasconcelos et al. (2021) tested the potential of DNA metabarcoding of bulk samples of young leaf tissues and successfully identified 34 species.

**Figure 6.**
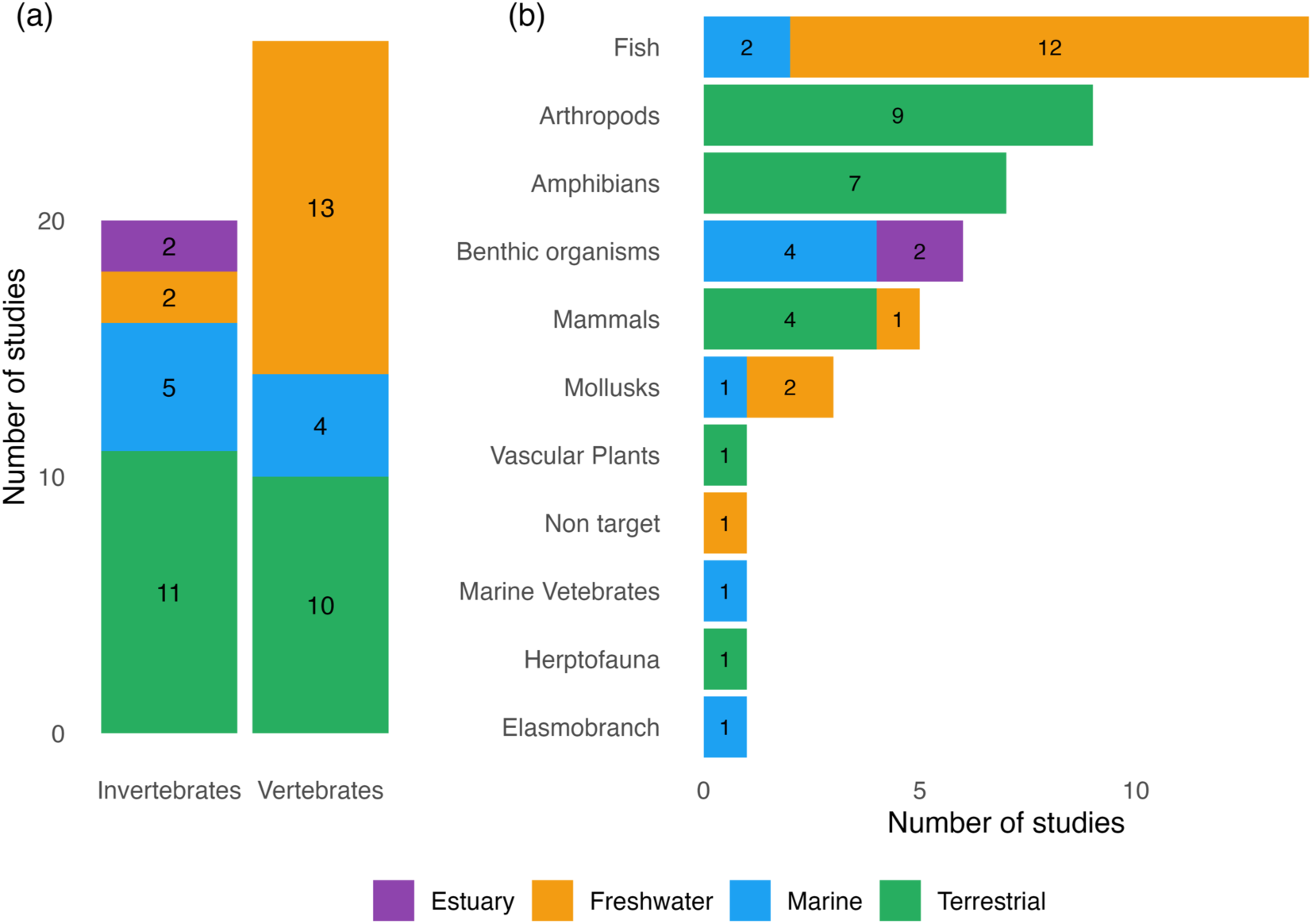
(a) Number of eDNA studies conducted in different ecosystems according to the target group (vertebrates or invertebrates). (b) Number of eDNA studies across ecosystems for each taxonomic group analyzed. Colors represent the ecosystem where each study was conducted.

Studies focusing on invertebrates encompassed both terrestrial and aquatic environments, targeting diverse taxa such as arthropods, mollusks, and benthic communities (Figure 6a, b). Within Arthropoda, investigations addressed groups like mosquitoes (family Culicidae), bees (clade Anthophila), and springtails (Collembola), the latter being soil-dwelling microarthropods. In the Cerrado biome, Martins et al., (2023) assessed the foraging behavior of Neotropical stingless bees through pollen and honey metabarcoding, an approach previously demonstrated only in Southeast Asia and Australia (Moura et al., 2022; Wilson et al., 2020). A key comparative study by de Faria et al. (2018) evaluated morphological and DNA metabarcoding methods in a subtropical bay, showing that DNA-based approaches retrieve a broader spectrum of marine meiofauna assemblages and enhance ecological analyses. Additional studies also used eDNA metabarcoding to characterize the composition and structure of marine meiofaunal (Coppo et al., 2024a, 2024b, 2025). Research on mollusks included a dietary analysis of *Doryteuthis sanpaulensis* using DNA metabarcoding (Ribas et al., 2021).

Among the studies investigating fish, two (14.3%) were conducted in marine environments and twelve (85.7%) in freshwater ecosystems (Figure 6b). Of these 14 fish studies, four freshwater studies and one marine study (de Queiroz et al., 2024) used DNA metabarcoding specifically focused on ichthyoplankton identification. The other study in marine environments targeting fishes aimed to analyze the diet of seven carangid species caught as bycatch in the Brazilian southwest Atlantic sardine fishery (Rosa et al., 2024). In freshwater ecosystems, studies focused mainly on fish biodiversity monitoring. Sales et al. (2021) applied eDNA metabarcoding to understand space-time dynamics in monitoring neotropical fish communities. One study aimed to understand the trophic interaction of vampire catfish through metabarcoding analysis of stomach content (Bonato et al., 2022). de Santana et al. (2021) evaluated eDNA metabarcoding as a complementary or alternative approach to traditional capture-based fish surveys in Amazonian rivers and streams. The method proved highly sensitive for detecting species and distinguishing fish communities across habitats, although species-level identification remains constrained by incomplete public DNA reference libraries in the region.

The potential of eDNA metabarcoding to detect vertebrates in aquatic environments was first explored in the Neotropics by Ritter et al. (2022). In their study, Ritter et al. (2022) noted that, unlike previous findings, the data mainly detected domestic and urban-associated species (cats, dogs, rats, cattle, ducks), with the capybara (*Hydrochoerus hydrochaeris*) as the only native non-domestic mammal. The authors suggest this method could support monitoring of common water-associated species and assess anthropogenic impacts through shifts from native to domestic fauna.

The seven studies involving amphibians identified in this review were all conducted in terrestrial environments (Figure 6b). Of these, five studies (71.4%) focused on the order Anura. Lopes et al. (2021) analyzed freshwater from tank bromeliads in the Atlantic Forest to detect eDNA of three threatened amphibians absent from traditional surveys. Although the targets were not found, DNA from other resident amphibians was successfully identified. Ernetti et al. (2024) reported the first potential detection of the invasive bullfrog *Aquarana catesbeiana* in southern Brazil’s Atlantic Forest. Sasso et al. (2017) showed that, despite some detection limitations, eDNA metabarcoding is an effective tool for assessing amphibian diversity in tropical streams and supporting Neotropical species conservation. In a related study, Lopes et al. (2019) detected eDNA from multiple eukaryotic taxa in leaf litter, including terrestrial amphibians, but found soil substrates less effective than water for amphibian detection. Lynggaard et al. (2019) used invertebrate-derived DNA (iDNA) and metabarcoding to demonstrate that vertebrate DNA can be detected through the analysis of bulk arthropod samples. A total of 32 vertebrate taxa were identified, including mammals, amphibians, and birds. The detected taxa were found within, or in proximity to, their known geographic distributions.

Regarding mammals, they were studied in four terrestrial research efforts (Figure 6b). Two of these focused on subterranean rodents (genus *Ctenomys*) in southern Brazil, specifically in the states of Santa Catarina and Rio Grande do Sul. In both studies, Lopes et al. (2015; 2020a) collected feces to perform diet analyses in order to understand ecological factors like niche overlap and resource competition between these co-occurring species. Lima et al. (2024) assessed the fauna of the Pantanal following the 2020 megafire by analyzing soil and water samples for mammal and herpetofauna detection. The study identified 27 mammal species across two sampling stations, while herpetofauna detections were comparatively low. Sales et al. (2019a) used water samples to evaluate terrestrial and aquatic mammal biodiversity in the Atlantic Forest and Amazon Rainforest. Across both biomes, representatives of eight mammalian orders and 14 families were detected. In the Amazon, six species were identified, three of which are listed as threatened by the IUCN (2019). In the Atlantic Forest, nine families were detected, with seven taxa identified at the species level. No study to date has targeted marine mammals (Figure 6b).

Regarding marine vertebrates, Lines et al. (2023) examined the lasting effects of the 2015 mining disaster on coastal biodiversity (Figure 6b). The authors collected saltwater samples near the mouths of the Rio Doce and Rio Jequitinhonha revealed that river discharge significantly influences coastal marine biodiversity, with strong interactions between rainfall and species richness in both basins. Seasonal shifts between tropical and subtropical water masses also play a key role in shaping biodiversity patterns in this ecoregion (Mazzucco and Bernardino, 2022).

### 3.5 Taxonomic units

Across all studies, OTUs represented the most frequently generated unit of taxonomic diversity (n = 24; 51%), followed by ASVs (n = 13; 31.9%). The combined use of both approaches was reported in two studies (4.3%), both conducted in freshwater environments targeting fish communities and using the 12S genetic marker (Figure 6a, b). We identified several studies that did not mention the generation of either ASVs or OTUs, but rather reported only the detected species (n = 10; 21.3%) (Figure 7a, b). Except for mammal and amphibian-focused studies, which consistently employed OTUs when generating taxonomic units or otherwise reported only species detections, the choice of taxonomic units varied among target organism groups (Figure 6b).

**Figure 7.**
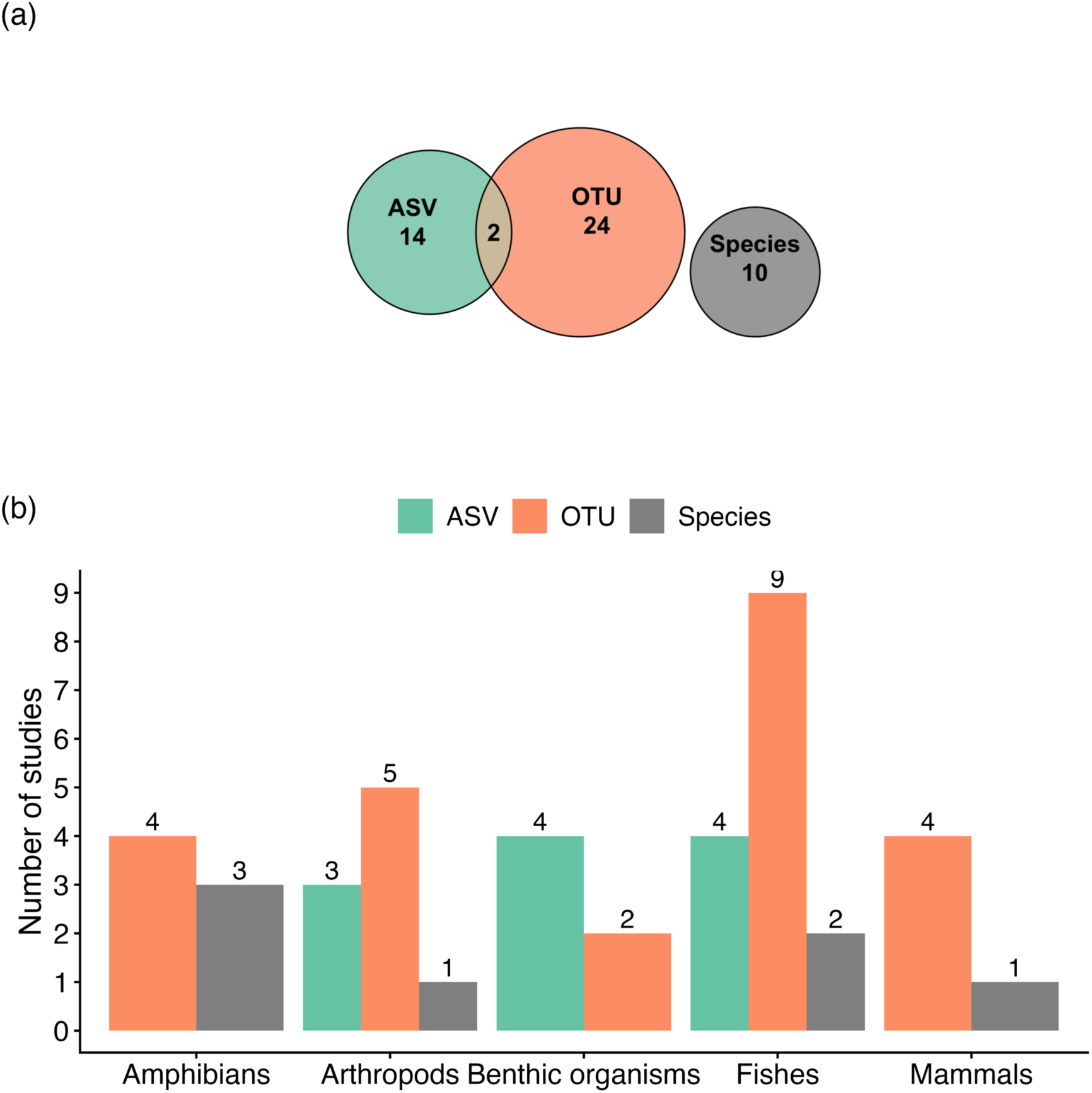
(a) Number of environmental DNA (eDNA) and/or metabarcoding studies that report different species units. (b) Number of studies reporting ASVs, OTUs, or Species across the six main target organisms in Brazil. ASV: amplicon sequence variant; OTU: operational taxonomic unit; Species: studies that reported only the detected species.

We did not identify a clear pattern in the taxonomic units used across ecosystems; however, in all four ecosystems assessed, both OTUs and ASVs were used. Hilário et al. (2023) demonstrated that both ASV and OTU-based measurements slightly overestimated species richness, yielding values higher than the actual number of species present in mock communities, as a single specimen often generated multiple ASVs/OTUs. Consequently, using ASVs or OTUs as direct proxies for species may lead to inflated estimates of biodiversity, with ASVs providing higher overestimation rates. For example, two studies conducted in estuarine habitats, both carried out in the same estuarine system, focusing on the same group of organisms and employing the same genetic marker, differed in the taxonomic units used. Bernardino et al. (2019) adopted ASVs, while Coppo et al. (2023) relied on OTUs. This divergence reflects a broader temporal trend, whereby ASVs have increased in prevalence from 2020 to 2023, despite OTUs remaining the predominant approach overall. This trend was also identified by Bunholi et al. (2023) in their systematic review.

Taxonomic units may include molecular operational taxonomic units (MOTUs), in many cases reported only as operational taxonomic units (OTUs). Traditionally, OTUs are generated by clustering sequence reads based on a high degree of nucleotide similarity, typically grouping reads that differ by less than 3%, corresponding to more than 97% sequence identity (Westcott and Schloss, 2015; Kopylova et al., 2016). In molecular marker data analysis, the creation of OTUs serves to reduce the impact of amplicon sequencing error by grouping errors with error-free sequences, and the observation of an OTU has been previously considered as a proxy of a ’species’ in taxonomic profiling (Callahan et al., 2017).

Alternatively, amplicon sequence variants (ASVs) differentiate sequences based on single-nucleotide resolution, inferring exact biological sequences in the sample by controlling for amplification and sequencing errors. ASVs are considered consistent labels with intrinsic biological meaning (Callahan et al., 2017; Forster et al., 2019). This provides significant advantages, including reusability across studies, reproducibility in future datasets, while also providing improved resolution and accuracy compared to traditional OTUs (Callahan et al., 2017; Forster et al., 2019). In this review, studies that reported the use of zero-radius OTUs (ZOTUs) were considered as applying ASVs approaches, given that both represent high-resolution sequence units generated by specific bioinformatic algorithms and aim to capture exact biological sequences at single-nucleotide resolution (Callahan et al., 2017; Edgar, 2017, 2018). The advent of DADA2 (Callahan et al., 2016), an open-source software tool facilitating the generation of ASVs, likely underpins this shift towards the adoption of ASVs in recent investigations (Callahan et al., 2017).

For conservation purposes, these findings underscore the need for caution when applying taxonomy-free metabarcoding, an approach increasingly proposed for investigating highly diverse and poorly characterized assemblages (Mächler et al., 2021), particularly in ecological assessments of the megadiverse regions. Despite certain limitations, the use of ASVs and OTUs remains effective for assessing species diversity and facilitates comparisons across multiple sites (Stahlhut et al., 2013) as well as spatiotemporal dynamics (Sales et al., 2021). In a recent meta-analysis, Zhang et al. (2025) reanalyzed 58 riverine fish eDNA datasets encompassing 1,818 sampling sites worldwide and found that species richness estimates and community structure metrics derived under a standardized bioinformatic workflow were generally consistent with those of the original analyses. However, the authors emphasized that differences in the reference database completeness directly affect the species identification accuracy.

### 3.6 Parachute Science

Among the 47 papers analyzed, 18 (38.3%) were conducted exclusively by Brazilian researchers and institutions. Three (6.4%) had foreign researchers as both first and senior authors, including foreign first authors’ nationalities from Denmark and Australia, with affiliations to the University of Copenhagen and Curtin University, respectively (Lynggaard et al., 2019; Lynggaard et al., 2020; Lines et al., 2023). Notably, all studies with a foreign first author also had a foreign senior author, and a total of eight (17%) papers had a foreign senior author only. Twelve studies (25.5%) received international financial support, and 10 of these (83.3%) were co-funded by Brazilian agencies (i.e., mixed financial support). Only two studies (4.3%) were funded exclusively by foreign institutions (e.g., United States Department of Agriculture - National Institute of Food and Agriculture (USDA-NIFA and University of Salford International Research Award), while 35 (74.5%) relied solely on Brazilian funding. Importantly, no study was conducted without the participation of at least one Brazilian author. It is important to highlight the potential risk of parachute science within eDNA and/or DNA metabarcoding applications, where research in low-income countries is carried out exclusively by international scientists from high-income countries, often without the involvement of local scientists or communities (Stefanoudis et al. 2021; Heyden, 2022). This may occur primarily due to insufficient research funding to support such costly investigations and the limited expertise available among local researchers in cutting-edge methodologies.

When considering the type of environment, six studies (30%) conducted in terrestrial ecosystems were carried out solely by Brazilian researchers. In marine ecosystems, four studies (44.4%) had only Brazilian representation, and six studies (37.5%) in freshwater systems were conducted without foreign collaboration. Of the two studies in estuarine habitats, only one involved exclusively local authors.

In some cases, the first author was a Brazilian national affiliated with a foreign institution. For classification, we prioritized nationality over institutional affiliation. In five studies (10.6%), first authors were Brazilians affiliated with foreign institutions such as the University of Duisburg-Essen (Germany) (Ritter et al., 2019), University of Salford (United Kingdom) (Sales et al., 2019a; Sales et al., 2019b; Sales et al., 2020) and the Zoological Research Museum Alexander Koenig (Germany) (Zenker et al., 2020). The countries most frequently involved in Brazilian eDNA and/or metabarcoding studies were the United States (n = 9; 19.1%) and the United Kingdom (n = 8; 17%), followed by Germany (n = 5; 10.6%), France (n = 4; 8.5%) and Australia (not considered Global South) (n = 3; 6.4%) (Figure 8). No country from the Global South had a significant participation. These patterns reflect a broader trend in the literature, the underrepresentation of Global South researchers as lead authors in studies conducted in their own regions, while Global North researchers are often prominently positioned, underscoring persistent structural inequalities in international collaborations (Jeffery, 2014; Stefanoudis et al., 2021; Miller et al., 2023).

**Figure 8:**
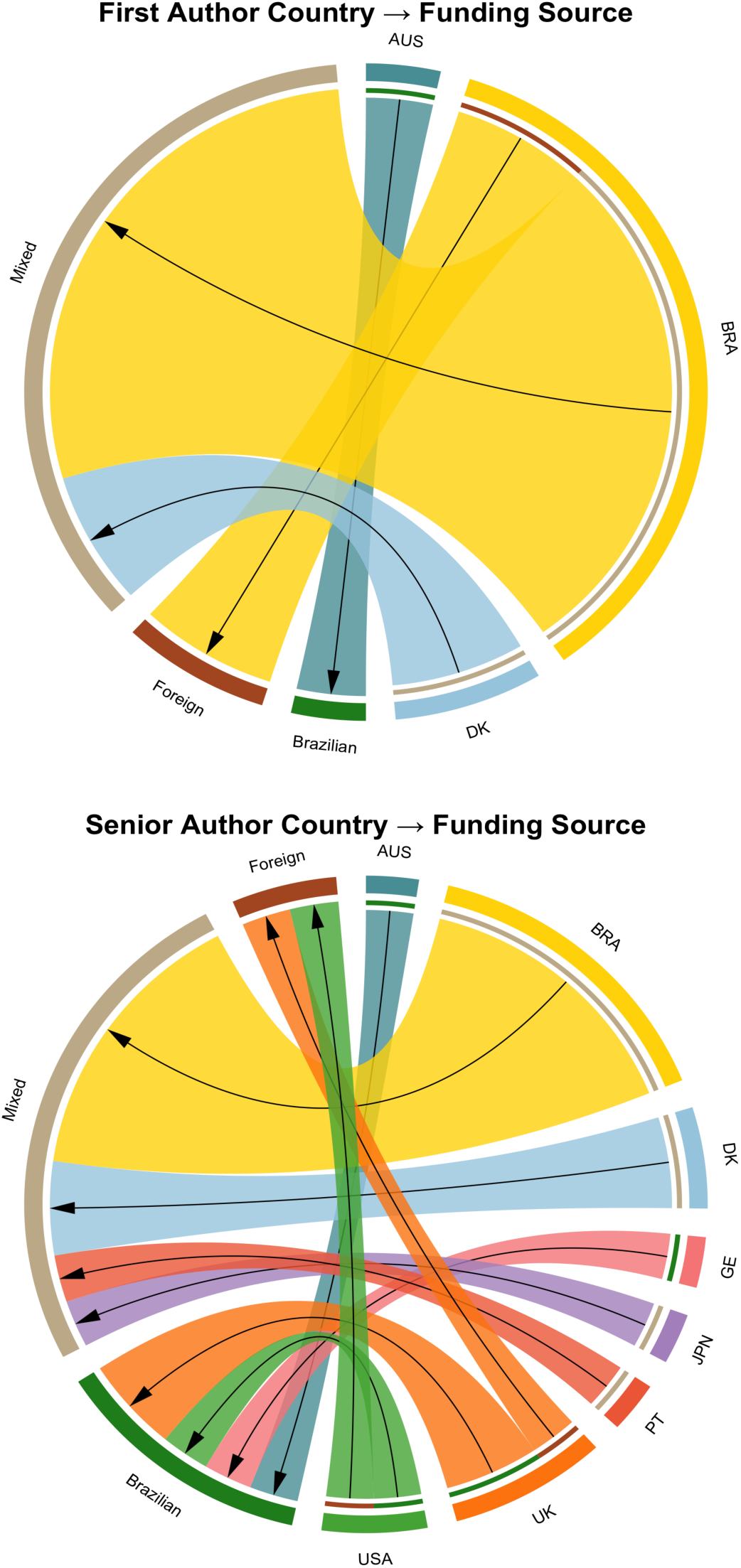
Chord diagrams showing the flows between author country affiliations and declared funding sources in environmental DNA (eDNA) and/or metabarcoding studies conducted in Brazil. First author affiliation; Senior author affiliation. Arrows connect the authors’ countries to the type of funding reported in each study (Foreign: funding received exclusively from foreign countries; Mixed: funding received by both Brazilian and foreign institutions; and Brazilian: funding received only by Brazilian institutions). To improve clarity, studies with exclusively Brazilian authorship and funding (i.e., studies in which both the first and last authors are affiliated in Brazil and received only Brazilian funding) were excluded from the figure. Country abbreviations: BRA: Brazil, USA: United States of America, UK: United Kingdom, GER: Germany, PT: Portugal, AUS: Australia, DK: Denmark, JPN: Japan.

Regarding financial support, the most frequently cited international agencies were the Agence Nationale de la Recherche (France), the Danish Council for Independent Research, and the Independent Research Fund Denmark (n = 2 each; 16.6% each). Broader initiatives also contributed, such as the CABANA Project (n = 2; 16.6%), a capacity-building initiative in Latin America funded by the Global Challenges Research Fund (UK Aid Budget), and the Global Genome Initiative (n = 1; 8.3%), which aims to preserve and share global genomic biodiversity (CABANA, 2025; Smithsonian Institution, 2025).

All studies funded exclusively by foreign institutions included at least one foreign first or senior author (Figure 8). Similarly, all studies with mixed (foreign + Brazilian) funding had foreign co-authors. Notably, foreign representation is higher among senior authors than first authors, which appears to influence the source of funding received (Figure 8). The institutions that funded the two exclusively foreign-financed studies were the USDA–National Institute of Food and Agriculture (USDA-NIFA) and the University of Salford. The most involved Brazilian agency in mixed funding scenarios was the National Council for Scientific and Technological Development (CNPq), appearing in 50% of these cases.

The first study using eDNA and/or metabarcoding in Brazil was published in 2015 and involved foreign funding and authorship (a French institution) (Lopes et al., 2015). In contrast, none of the 2 studies published in 2025 featured foreign authors. From 2024 to the present, eight studies were published, with only two featuring any foreign co-authors, both from institutions in the United States.

This review revealed only weak signs of parachute science in Brazilian eDNA and metabarcoding studies, since no studies were conducted without a Brazilian representation, and 38.3% of the studies were conducted exclusively by Brazilian researchers and institutions. However, Brazil continues to face challenges common to many Global South countries, including limited access to research funding and equipment (Trisos et al., 2021). More importantly, Brazil lacks specialized infrastructure for this type of study, which requires dedicated rooms for eDNA extraction as well as separate rooms for pre- and post-PCR procedures. This setup helps prevent different types of contamination and ensures the strict quality control required for this kind of research.

To sustain the observed reduction in parachute science and foster more equitable and inclusive global research, it is essential to support local leadership, improve access to funding and data, and promote collaborative models that prioritize capacity-building in the Global South.

### 3.7 Challenges and future perspectives for the use of eDNA and DNA metabarcoding for biodiversity conservation

A major challenge identified in Brazilian DNA metabarcoding and eDNA studies is the lack of comprehensive and accessible DNA reference libraries, a limitation shared worldwide (Ruppert et al., 2019; Jackman et al., 2021; Zainal et al., 2022; Morey et al., 2024). Expanding these libraries through natural history collections and vouchered specimens, along with investments in training and outreach, is essential for improving biodiversity assessment (de Santana et al., 2021). Despite gaps in reference data, eDNA has proven to be a reliable and cost-effective biomonitoring tool in the Neotropics (Dal Pont et al., 2021). Regional barcode databases can further enhance restoration and vegetation surveys (Vasconcelos et al., 2023), while custom reference sets and controlled experiments are key to increasing taxonomic resolution and methodological accuracy (Hilário et al., 2023).

In order to overcome this challenge, over the past decade, various initiatives have launched efforts to sequence reference-level genomes and implement large-scale monitoring using eDNA (Vilaça et al., 2025). In Brazil, a major national initiative called Genomics of the Brazilian Biodiversity (GBB) consortium, which prioritizes the generation of high-quality genomic data, including barcode references and eDNA to support conservation and unlock bioeconomic potential (Vilaça et al., 2025). These efforts are supported by investments in local sequencing infrastructure and a strong focus on capacity building and collaboration, aiming to overcome historical asymmetries and ensure that knowledge and data remain within the country (Vilaça et al., 2025)

Beyond the technical limitations, structural and historical barriers also play a role. Building open-access data platforms can provide a solid foundation for truly globalizing conservation science. In addition, there is a clear need to strengthen the national infrastructure for eDNA research. Like most emergent and developing nations, Brazil experiences high operating costs for scientific laboratories because most reagents and equipment are imported from the Global North (Helmy et al., 2016). Considering the vast geographical area of the country, a limited number of laboratories in Brazil are fully equipped and experienced in handling environmental DNA workflows, from contamination-free sampling and molecular processing to the complexities of bioinformatic analysis. This limitation hampers the scalability and reliability of eDNA-based monitoring, especially in regions outside major research centers. Notably, no studies involving eRNA were identified in our initial survey of Brazilian publications, highlighting this as a potential area for future exploration.

The availability of public funding is another critical constraint. Investments in biodiversity monitoring remain low compared to the urgency of environmental degradation in the country. A preliminary search in major Brazilian funding agencies, such as CNPq and FAPESP (São Paulo Research Foundation), reveals a relatively small number of active research projects specifically focused on eDNA and metabarcoding. As of June 2025, the São Paulo Research Foundation (FAPESP) supports only six active research grants, ten ongoing national scholarships, and one international scholarship related to environmental DNA, totaling 17 ongoing initiatives (FAPESP, 2025). Expanding these funding lines, particularly for interdisciplinary, applied, and long-term research, will be crucial to ensure the continuity, innovation, and consolidation of this field in Brazil. Equally important is the capacity-building of specialized personnel. Training researchers and technicians in all stages of eDNA analysis, such as field sampling, molecular techniques, and data interpretation, continues to be of urgent importance. Most current initiatives rely on collaboration with foreign laboratories or short-term training programs, which are often insufficient to foster long-term autonomy and technical excellence.

Finally, conservation and ecological understanding have benefited from DNA metabarcoding and eDNA studies. These DNA-based methods contribute not only to species identification but also to addressing several key issues relevant to conservation. Although species identification remains a cornerstone of effective management, these genetic approaches extend their conservation applications to areas such as detecting and assessing bioinvasions, ecosystem-based biomonitoring, evaluating the effectiveness of marine protected areas, and supporting sustainable fisheries management. However, realizing the full potential of eDNA and metabarcoding for biodiversity monitoring requires overcoming both technical and systemic challenges. These include improving reference databases through genome and mitogenome sequencing, incorporating advanced computational tools, investing in capacity-building, promoting equitable access to data and biological collections, and strengthening local scientific infrastructure. Addressing these challenges is essential for the effective application of eDNA and metabarcoding in conservation efforts in Brazil and other biodiversity-rich regions of the Global South.

## ACKNOWLEDGMENTS

This work was supported by the São Paulo Research Foundation (FAPESP #2019/10201-0 and #2024/16716-0)

